# Local coupling between sleep spindles and slow oscillations supports the stabilization of motor memories

**DOI:** 10.1101/2020.08.24.264697

**Authors:** Agustín Solano, Luis A. Riquelme, Daniel Perez-Chada, Valeria Della-Maggiore

**Affiliations:** IFIBIO Houssay; Department of Physiology, School of Medicine, University of Buenos Aires, Argentina; Department of Internal Medicine, Pulmonary and Sleep Medicine Service, Austral University Hospital, Buenos Aires, Argentina

## Abstract

Recent studies from us and others suggest that traditionally declarative structures (e.g., hippocampus) mediate some aspects of the encoding and consolidation of procedural memories. This evidence points to the existence of converging physiological pathways across memory systems. Here, we examined whether the coupling between slow oscillations (SO) and spindles, a mechanism well established in the consolidation of declarative memories, is relevant for the stabilization of human motor memories. To this aim, we conducted an EEG study in which we quantified various parameters of these oscillations during a night of sleep that took place immediately after learning a visuomotor adaptation task. We hypothesized that if this coupling is instrumental to motor memory consolidation then spindles locked to the active phase of a slow oscillation would predict long-term memory. We found that visuomotor adaptation increased the overall density of fast (≥12 Hz) but not slow (<12Hz) spindles during NREM3. This modulation was manifested rather locally, over the hemisphere contralateral to the trained hand. Although motor learning did not affect the density of SOs, it substantially enhanced the number of fast spindles locked to the active phase of SOs. The fact that only coupled spindles of the left hemisphere predicted long-term memory overnight, points to the precise phase relationship between these oscillations as a fundamental signature of motor memory consolidation. Our work provides evidence in favor of a common mechanism at the basis of the stabilization of declarative and non-declarative memories.

**Significance Statement:** Ever since the discovery of memory systems, the study of the mechanisms supporting the consolidation of declarative and procedural memories has progressed somewhat in parallel. In the last few years, however, this framework is starting to change. We now know that structures originally thought of as purely declarative, such as the hippocampus, participate in the consolidation of procedural tasks. Here, we show that sleep modulates the stabilization of motor memories through a mechanism involved in the consolidation of declarative memories, based on the local synchrony between fast sleep spindles and slow oscillations. The fact that only coupled –but not uncoupled- spindles of the contralateral hemisphere predicted long-term memory supports a role of this association in the consolidation of motor memories.

## Introduction

In the last decades, the function of sleep in memory consolidation has received extraordinary attention. There is now compelling evidence from human and non-human studies supporting a role of NREM sleep in both the stabilization and enhancement of declarative memory (e.g., Plihal & Born, 1997; Rasch et al., 2007; Diekelmann & Born, 2010; Schönauer et al., 2015). The neural signatures of sleep dependent consolidation are rapidly being unraveled. We now know that during NREM sleep slow oscillations (SO, ~1 Hz) generated in the cortex (Amzica & Steriade, 1998; Timofeev et al., 2000) facilitate the occurrence of thalamic spindles (~ 10-16 Hz) during their excitable -up-state- (Steriade, 2006; Staresina et al., 2015; Schönauer & Pöhlchen, 2018). A recent study has in fact shown that the induction of thalamic spindles through optogenetics potentiates hippocampus-dependent memories when stimulation is phase locked with the up-state (active phase) of SOs, whereas spindle suppression impairs memory formation (Latchoumane et al., 2017). It is the precise synchrony between *fast* spindles (≥12 Hz) and SOs, in conjunction with hippocampal sharp-wave ripples, what appears to mediate systems consolidation in a variety of declarative tasks (Buzsaki, 2015; Maingret et al., 2016; Ladenbauer et al., 2017; Helfrich et al., 2018; Muehlroth et al., 2019; Navarro-Lobato & Genzel, 2019).

Much less is known about the neural signatures of motor memory consolidation during sleep. Motor learning comprehends motor skill learning (MSL) and motor adaptation (MA). The first involves the acquisition of new motor plans whereas the second, the recalibration of existing motor plans (Krakauer el al., 2019). Growing evidence suggests that offline gains observed in humans during motor sequence learning, a type of MSL, are linked to the hippocampus (Albouy et al., 2013b; Albouy et al., 2015; Döhring et al., 2017; Schapiro et al., 2019; Jacobacci et al., 2020), and relate to an increment in the density of sleep spindles (Nishida & Walker, 2007; Morin et al., 2008; Barakat et al., 2011; Albouy et al., 2013a; Boutin et al., 2018). Recent work carried out in rats indicates that learning an MSL task is associated with neural replay over the motor cortex, often phased locked to a SO (Ramanathan et al., 2015), and with an enhancement of the coupling between spindles (fast & slow) and SOs (Silversmith et al., 2020). Yet, whether these changes impact on overnight improvements in performance has not been explored. Finally, a couple of studies have shown that MA increases the power of slow -delta- waves (~1-4 Hz) during the early portion of NREM sleep that predicts overnight improvements in performance (Huber et al., 2004; Landsness et al., 2009). Yet, whether the synchrony between SO and spindles is relevant for the consolidation of adaptation memories remains unknown.

In this study, we examined whether the coupling between spindles and slow oscillations supports the stabilization of visuomotor adaptation (VMA) memories. Given the role of fast spindles in the consolidation of declarative memories we also explored the contribution of fast and slow spindles to this process. We have previously shown behavioral evidence supporting the diurnal consolidation of VMA memories within a 6-hour window (Albert et al., 2019; Lerner et al., 2020). We have also reported that, during this period, the increment in the functional connectivity between motor and parietal areas contralateral to the trained hand predicts long-term memory (Della-Maggiore et al., 2017). Here, we addressed whether these local changes in brain activity are further modulated by sleep. To this aim, we took advantage of a protocol focused on the close proximity between learning and sleep, known to improve overnight memory retention in declarative and motor tasks (e.g., Huber et al., 2004; Landsness et al., 2009; Schönauer et al., 2015). After a night of familiarization, volunteers performed a visuomotor adaptation session or a control session and we quantified the density of spindles (fast and slow), and the coupling between spindles and slow oscillations during NREM sleep. We predicted that if the level of coupling between these oscillations is involved in the stabilization of VMA memories, then it should increase locally and predict long-term memory overnight.

## Materials and Methods

### Participants

Ten normal volunteers (5 females, age: (mean ± SD) 24.3±3.1 years old) were recruited for this study. All subjects were right-handed as assessed by the Edinburgh Laterality Questionnaire (Oldfield, 1971), and none of them declared neurological nor psychiatric disorders. All potential participants were asked to fill the Pittsburgh Sleep Quality Questionnaire (Buysse et al., 1989) and were evaluated on the Epworth Drowsiness Scale (Johns, 1991). Only subjects fulfilling the criteria for good sleep quality based on the last two indices were included in the study. Participants were instructed to maintain a regular sleep schedule and to avoid drinking alcohol and coffee during the duration of the experiment. This was monitored through self-recorded spreadsheets provided by the researcher.

All volunteers signed the informed consent approved by the Ethics Committee of the Hospital de Clínicas, University of Buenos Aires (approved on Nov, 24, 2015, and renewed every year), which complies with the Declaration of Helsinki in its latest version (Fortaleza, 2013), and with the National Law on the Protection of Personal Data.

### Experimental paradigm

The visuomotor adaptation (VMA) task has been described in detail elsewhere (Villalta et al., 2015; Lerner et al., 2020) and will be briefly summarized here. Subjects performed a center-out task consisting of moving a cursor from a start point in the center of a computer screen to one of 8 visual targets arranged concentrically 45° from each other using a joystick, controlled with the thumb and index finger of the right hand (Figure 1.A). Vision of the hand was occluded. Participants were instructed to make a shooting movement to the target. One cycle consisted of eight trials, in which subjects made pointing movements to each of the eight targets, whereas there were 11 cycles per block.

**Figure 1.**
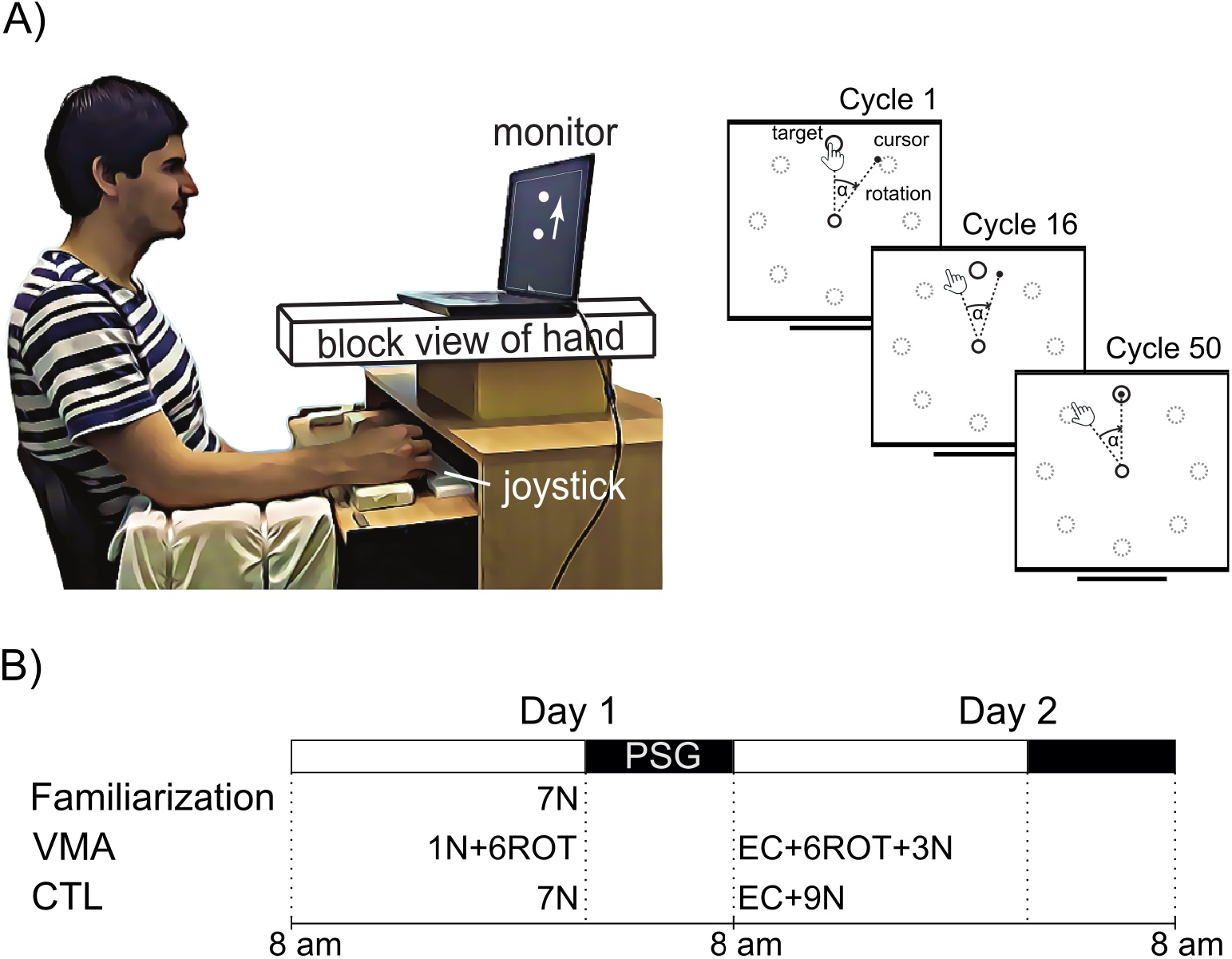
Experimental setup and experimental design. *A) Experimental setup.* Subjects sat on a chair and performed center-out movements to one of 8 visual targets using a joystick controlled with their right hand (Left panel). The right panel is a cartoon representing the visual display and illustrates the effect of applying the visuomotor rotation (ɑ) over the movement direction of the hand and the cursor as a function of training (cycles). As adaptation progressed over cycles of practice, subjects learned to bring the cursor to the target. *B) Experimental design.* All subjects went through an initial session of familiarization, followed by a visuomotor adaptation session (VMA) and a control session (CTL), each separated by 1 week. The order of VMA and CTL sessions was counterbalanced. During the familiarization session they performed 7 blocks of null trials (N). During the VMA session, they performed 1 block of null trials followed by 6 blocks of perturbed trials in which a visual rotation was applied (ROT). On Day2, retention was assessed using 2 error-clamp cycles (EC), and then they re-adapted to the same perturbation (6ROT), and were subsequently washed out (3N). During the CTL session, subjects performed the corresponding number of blocks without the visual rotation (N). Sleep was monitored with polysomnography (PSG) overnight, starting ~10 minutes after performing the task on Day 1.

Subjects performed three different types of trials throughout the study. During perturbed trials a clockwise 45-degree visual rotation was imposed to the cursor relative to the movement of the hand (see Figure 1.A, right panel). During null, i.e., unperturbed trials in which no visual rotation was applied, the movement of the cursor directly mapped onto the hand movement. Finally, during error-clamp (EC) trials, visual feedback was manipulated to provide fake “straight” paths to the target (mean error=0°, SD=10°). Error-clamp trials were used to assess memory retention, and served to estimate the state of the motor system while preventing participants from continuing to learn from error (Criscimagna-Hemminger & Shadmehr, 2008). The VMA task was implemented in MATLAB (The MathWorks, Inc, MA) using the Psychophysics Toolbox v3 (Brainard, 1997; Kleiner et al., 2007).

### Experimental Design

To examine whether the coupling between sleep spindles and slow oscillations relates to the consolidation of motor memories we conducted a longitudinal experiment following a within-subjects repeated-measures design, consisting of three sessions separated by 7 days each: i) a familiarization session during which subjects performed the center-out task during null trials (N), that is, in the absence of the visual rotation, ii) a visuomotor adaptation session (VMA) in which subjects adapted to a visual rotation (ROT), and iii) a control session (CTL) in which subjects performed the center-out task in the absence of the optical rotation (N). We chose a repeated measures design to reduce inter-individual variance and improve statistical power. The experimental design is described in Figure 1.B. After the familiarization session, subjects were randomly assigned to the VMA or the CTL condition, each of which took place on a different day and week. The order was counterbalanced so that half of the volunteers performed the VMA condition first and the other half second. The familiarization session was included for two reasons. First, given that subjects have an internal bias for movement direction, adaptation may even take place in the control condition (e.g. Della-Maggiore et al., 2017). The familiarization session thus served as a first approximation to practice with the joystick and the experimental paradigm in the absence of the optical rotation. Second, Tamaki and collaborators (2016) have shown that subjects do not sleep well during the first night in a new environment. Familiarization with the experimental setup thus aimed at improving the PSG recordings.

Participants arrived to the sleep laboratory at 9 pm and electroencephalography (EEG) electrodes were placed over their scalp following the 10-20 montage (see below). Next, they performed the center-out task and went to bed ~10 min later for a full night of sleep. A PSG recording was obtained throughout the night. During the familiarization session, participants performed seven blocks of the task without any perturbation (7N). During the VMA and CTL sessions, subjects performed the task before (Day 1) and after (Day 2) a full night of sleep. In the VMA session, on Day 1, they performed one block of null trials (1N) followed by six blocks of perturbed trials in which the visuomotor rotation was applied (6ROT). On Day 2, they performed two cycles of error-clamp trials (EC) to assess retention, followed by 6 blocks of perturbed trials (6ROT) and three blocks of null trials (3N) to washout. The CTL session followed the same experimental protocol but the visual rotation was not applied (7N on Day 1, followed by EC+9N on Day 2). In all cases, the level of vigilance before performing the task was assessed using the Stanford Sleepiness Scale (Hoddes E., 1973).

### Polysomnographic recordings (PSG)

Eleven surface electroencephalogram (EEG) electrodes were placed over prefrontal, motor and parietal areas (FC1, FC2, FC5, FC6, C3, C4, P3, P4), and over the midline (Fz, Cz, Pz). Electrodes were mounted following the standard 10-20 arrangement (Modified Combinatorial Nomenclature; Oostenveld & Praamstra, 2001). Both mastoids were used as references. In addition to EEG electrodes, two electrodes were placed over the periorbital area of both eyes and two additional electrodes over the chin, to measure electrooculography (EOG) and electromyography (EMG), respectively. All signals were acquired at 200 Hz, using the Alice 5 EEG equipment (Philips Respironics, PA, EEUU).

### EEG processing

EEG signal was bandpass-filtered between 0.5 and 30 Hz using a FIR filter (*eegfiltnew* function) from the MATLAB's toolbox EEGLAB (Delorme & Makeig, 2004), and processed with the Artifact Subspace Reconstruction algorithm (ASR, Mullen et al., 2015) to remove transient and large-amplitude artifacts (*clean_rawdata* function from EEGLAB; threshold= 30). EOG and EMG signals were also bandpass-filtered in order to facilitate sleep scoring (filter cutoff frequencies: EOG = 0.5-15 Hz; EMG = 20-99 Hz). All PSG recordings were sleep staged manually, according to standard criteria of the American Academy of Sleep Medicine (AASM; Iber, 2004). Namely, 30-second epochs were classified as either Wake (W), Non Rapid Eye Movement (NREM1, NREM2 and NREM3), or Rapid Eye Movement (REM) stage. Stage classification was carried out by two independent scorers and a concordance criterion was used to define the stage of each epoch. After stage classification, sleep architecture was determined based on the metrics mentioned below (Data Analysis). Movement artifacts on the filtered EEG signal were detected by visual inspection and manually rejected.

Slow oscillations (0.5-1.25 Hz) and sleep spindles (10-16 Hz) were automatically identified from the EEG signal corresponding to the stages NREM2 and NREM3, using previously reported algorithms (see below).

#### Detection of Slow oscillations (SO)

The algorithm implemented to detect SOs was based on that reported by Mölle and collaborators (2011) and Antony & Paller (2016). EEG signal was bandpass-filtered between 0.5 and 1.25 Hz. To quantify SOs, we first identified zero crossings of the EEG signal and labeled them as positive-to-negative (PN) or negative-to-positive (NP). Next, those EEG segments between two NP zero crossings were considered a slow oscillation if i) they lasted between 0.8 and 2 seconds, ii) their maximum amplitude was positive, and iii) their minimum amplitude was negative. This algorithm was used for each channel of each session. Only slow oscillations with a peak-to-peak amplitude greater than the median amplitude of the corresponding channel were kept for subsequent analysis (Mizrahi-Kliger et al., 2018).

#### Sleep spindles detection

The algorithm implemented to detect sleep spindles was based on that reported by Ferrarelli and collaborators (2007) and Mölle and collaborators (2011). It was run along each channel for each session. First, EEG signal was bandpass-filtered between 10 and 16 Hz before calculating the instantaneous amplitude (IA) and instantaneous frequency (IF) by applying the Hilbert Transform (Tort et al., 2010). The IA was used as a new time series and smoothed with a 350 ms moving average window. Next, those segments of the IA signal that exceeded an upper magnitude threshold (90^th^ percentile of all IA points) were labeled as potential spindles. The beginning and end of potential spindles were defined as the points in which the signal dropped below a lower threshold (70^th^ percentile of all IA points). Potential spindles with a duration between 0.5 and 3 seconds were labeled as true spindles; mean frequency, duration and maximum peak-to-peak amplitude were calculated for each true spindle. Finally, spindles were further classified into two types according to their mean frequency: slow spindles with a frequency < 12 Hz and fast spindles with a frequency ≥ 12 Hz (Mölle et al., 2011; Cox et al., 2017).

#### Coupling between slow oscillations and spindles

After identifying spindles and SOs, we looked for spindles that occurred during a SO. We quantified spindle-SO couplings according to the following criterion: if a spindle had its maximum peak-to-peak amplitude during the course of a SO it was counted as a spindle-SO coupling.

In order to further characterize the level of association between these oscillations we explored the phase of the SO at which the spindle occurred using the method reported by Tort et al. (2010) and Niknazar et al. (2015). To this end, the instantaneous phase (IP) of each SO was first obtained through the Hilbert transform. The IP of an oscillation varies in a range of ±π radians, so we arbitrarily fixed the SO’s positive peak to 0 radian and the negative peak to either + π or −π radians (PN and NP zero crossings occurred at +π/2 and −π/2 radians, respectively). The phase relationship for each spindle-SO coupling was defined as the phase of the SO at which the spindle developed its maximum peak-to-peak amplitude. This algorithm was applied to each channel of each session.

### Data analysis

#### Behavior

Motor performance was measured based on the movement direction of the joystick (pointing angle) relative to the line segment connecting the start point and target position. Trials in which the pointing angle exceeded 120 degrees were excluded from further analysis. Given that trials were organized in cycles of eight targets, all trial-by-trial data were converted into cycle-by-cycle time series by computing the median pointing angle for each cycle and each subject. To assess memory retention, the median of error-clamp cycles obtained overnight was computed and expressed as a percentage of the median corresponding to the last block of learning (asymptote).

#### EEG signal

##### - Sleep architecture

Sleep architecture was determined based on the following measures expressed in minutes: total sleep time, sleep latency (latency to NREM1), REM latency, total wake time, WASO (wake after sleep onset), time in NREM1, NREM2, NREM3 and in REM. Sleep efficiency was also computed as the percentage of total sleep time relative to the time interval between lights-off and lights-on (%).

##### - SO and Spindle measures

As described above, SOs and spindles were automatically identified from the EEG signal previously classified as NREM2 and NREM3. Given that the EEG signal from some channels deteriorated due to friction as the night progressed, we focused the EEG analysis on the first sleep cycle. The following measures were computed for sleep spindles: duration (msec), mean frequency (Hz), peak-to-peak amplitude (μV) and density of fast and slow spindles (number of sleep spindles per minute of NREM2 or NREM3 sleep). Density was also computed for SOs (number of SOs per minute of NREM3 sleep). To characterize the spindle-SO coupling we computed the proportion of coupled spindles, the density of spindles coupled with a SO (number of spindle-SO couplings per minute of NREM3 sleep), and their coupling phase.

To assess how these measures differed between VMA and CTL we computed their relative difference according to the function ((VMA-CTL)/CTL*100) for each EEG channel and each subject. We first conducted a global statistical analysis on this measure across the 11 electrodes for each of the sleep metrics of interest. To appreciate the spatial distribution of the effects we report the results for VMA, CTL and their relative difference in topographic maps (MNE-Python; Gramfort et al., 2013). Furthermore, to explore the possibility of a local – hemispheric- modulation (Nishida & Walker, 2007; Johnson et al., 2012; Della-Maggiore et al., 2017; Geva-Sagiv & Nir, 2019), we also conducted separate statistical analyses on the data pooled across the channels of the left hemisphere (LH: electrodes FC1, FC5, C3 & P3) and right hemisphere (RH: electrodes FC2, FC6, C4 & P4). Given that the midline may capture electrical activity from both hemispheres it was not included in the latter analysis.

In order to study the level of synchrony between fast spindles and SOs we determined the phase of the SO at which the spindle yielded its maximum peak-to-peak amplitude and contrasted the level of grouping of spindles around the mean phase for the VMA and CTL sessions. Specifically, we first computed the mean coupling phase for each subject and across subjects in polar coordinates, and displayed them in circular plots. The level of synchrony was then assessed by comparing the variance of the individual coupling phase around the mean across sessions.

To visually illustrate the synchrony and phase relationship between slow oscillations and spindles, the grand average of the EEG signal filtered for the slow oscillations frequency band (0.5-1.25 Hz) and the grand average of the *fast* sleep spindles frequency band (12-16Hz) were graphically overlaid, time locked to the spindle’s maximum amplitude.

### Statistical analysis

Statistical analyses were carried out using R (version 3.4.1; R Core Team, 2017) in RStudio (Rstudio Team, 2015). Statistical differences were assessed at the 95% level of confidence. When the variables of interest followed normality and homogeneity of variance, statistics were carried out by fitting a Linear Mixed Model (LMM, using the ‘lmer’ function implemented in the ‘lme4’ package in R, Bates et al., 2015). Random intercepts of LMM were estimated for each subject to take into account the repeated measures across sessions. The response variable was either the relative difference between sessions ((VMA-CTL)/CTL*100) or the measures corresponding to the VMA and CTL conditions. The fixed effects were the condition (VMA or CTL), sleep stage (NREM2 or NREM3), the type of spindle (fast or slow), the hemisphere (left or right hemisphere) and the sleep stage by spindle type interaction, depending on each analysis. To assess the statistical significance of fixed effects, we used a step-up strategy starting with an empty model (global mean of the response variable) and then we successively added potential predictors (fixed effects). We used the *Likelihood Ratio Test* to compare the fitted models (‘anova’ function in R). One sample t-tests were used to perform global statistical comparison versus zero for the relative difference between sessions.

In cases where the statistical distributions failed tests for normality or homogeneity of variance data was analyzed using non-parametric tests. For repeated measures, the Friedman test was used in the case of having three or more conditions.

In cases where two circular distributions were compared, we used the Wallraff’s Nonparametric Test for circular concentration (homoscedasticity). The Rayleigh test was used to assess unimodal departure from uniformity for each circular distribution. All tests are implemented in the ‘circular’ R package (Agostinelli & Lund, 2017).

Finally, to examine if the density of SO-coupled and uncoupled spindles was associated with long-term memory overnight, a Pearson correlation was computed for each measure and corrected for multiple comparison using the Bonferroni correction (adjusted alpha level = 0.025).

## Results

To examine whether the coupling between sleep spindles and slow oscillations may contribute to the stabilization of visuomotor adaptation memories, we quantified these measures during the period of NREM of the first sleep cycle. We also examined various metrics of the sleep architecture to assess whether learning affected any sleep stage/s differentially. A group of subjects went through an initial familiarization session followed by either a session of visuomotor adaptation (VMA), in which subjects compensated for a visual rotation, or a control session (CTL), in which the visual rotation was not applied (Figure 1.B). The three sessions were separated by a week. During the VMA and CTL sessions participants performed the motor task both before and after a full night of sleep. Memory retention was assessed overnight before the second practice session using error-clamp trials (EC).

### Behavioral Results

All volunteers learned to compensate the visual rotation during the VMA session and, on average, retained 56±10.5 % (mean ± SE) overnight. Moreover, the degree of vigilance assessed using the Stanford Sleepiness Scale yielded scores below 3 (Day 1 and Day 2 for CTL and VMA sessions = p>0.5), consistent with a mental state of alertness both before performing the task at night and the day after. For further behavioral details on these measures, refer to the Extended Data (Figure 1-1 and Table 1-1, respectively).

**Table 1.** Sleep Architecture. Shown are the mean and standard deviation (SD) corresponding to the sleep measures listed in the first column for all conditions. The last column depicts the statistics and p values yielded by statistically comparing the three conditions. All measures are depicted in minutes except for Sleep Efficiency, defined as the percentage of total sleep time relative to the time interval between lights-off and lights-on (%). WASO: wake after sleep onset.

| Measure | Familiarization |  | CTL |  | VMA |  | Friedman test |  |
| --- | --- | --- | --- | --- | --- | --- | --- | --- |
|  | Mean | SD | Mean | SD | Mean | SD | Chi-sq | p |
| Total Sleep Time (min) | 376.40 | 75.81 | 386.10 | 28.20 | 386.85 | 43.52 | 2.60 | 0.27 |
| Sleep Efficiency (%) | 81.90 | 16.69 | 87.36 | 6.92 | 86.57 | 9.86 | 0.80 | 0.67 |
| Sleep latency (min) | 18.20 | 15.66 | 18.40 | 13.75 | 16.05 | 10.75 | 0.20 | 0.90 |
| REM latency (min) | 132.60 | 65.80 | 105.00 | 31.27 | 123.85 | 49.53 | 0.60 | 0.74 |
| Total Wake Time (min) | 80.35 | 77.47 | 54.80 | 30.59 | 58.75 | 45.80 | 0.20 | 0.90 |
| WASO (min) | 42.95 | 63.03 | 33.10 | 28.29 | 29.50 | 20.24 | 1.08 | 0.58 |
| NREM1 (min) | 43.35 | 23.51 | 53.30 | 30.63 | 53.35 | 16.27 | 0.60 | 0.74 |
| NREM2 (min) | 157.30 | 55.89 | 157.25 | 36.30 | 150.15 | 46.48 | 0.60 | 0.74 |
| NREM3 (min) | 100.25 | 38.80 | 100.15 | 23.56 | 105.90 | 23.49 | 1.40 | 0.50 |
| REM (min) | 75.50 | 34.54 | 75.40 | 29.18 | 77.45 | 26.84 | 0.60 | 0.74 |

### Effect of VMA on sleep architecture

We found no significant differences in sleep architecture across sessions for any of the computed measures, suggesting that adaptation did not modulate the intrinsic sleep structure. All scores and corresponding statistics are displayed in Table 1.

### Visuomotor adaptation modulates the density of fast spindles during NREM3

We first looked at the impact of visuomotor adaptation on the intrinsic features of spindles, that is, frequency, duration and peak-to-peak amplitude across NREM sleep. No statistical differences were found for frequency (LMM stats, main effect of session; χ^2^(1)=0.07, p=0.79), duration (LMM stats, main effect of session; χ^2^(1)=0.46, p=0.49), or amplitude of sleep spindles (LMM stats, main effect of session; χ^2^(1)=0.03, p=0.87).

Next, we examined the global effect of motor learning on spindle density during NREM sleep. Previous work has distinguished between two types of spindles according to their spatial distribution and intrinsic frequency (Mölle et al., 2011; Cox et al., 2017). Fast spindles, which have been linked to memory consolidation (Barakat et. al., 2011; Ladenbauer et al., 2017; Helfrich et al., 2018; Muehlroth et al., 2019; Navarro-Lobato & Genzel, 2019), are distributed over centro-parietal areas and have a frequency ≥12 Hz, whereas slow spindles are distributed over frontal areas and have a frequency <12 Hz. Here, we explored whether VMA differentially modulated fast and slow spindles. In humans, motor sequence learning has been associated with an increment in the density of fast spindles during NREM2 (e.g. Boutin et al., 2018). Given that we were interested in studying both the number of spindles and their coupling with slow oscillations, which are more prominent during NREM3, we quantified spindle density for both sleep stages.

Figure 2 depicts the effect of VMA on the density of fast and slow spindles during NREM2 and NREM3 (relative change in density, mean ± SE: *NREM2: Fast spindles=−1.33±3.76 %, Slow spindles=−3.39±3.28 %; NREM3: Fast spindles=22.59±4.21, Slow spindles=−3.42±1.59 %*). We found that adaptation to a visual rotation increased the density of sleep spindles during NREM3 but not during NREM2 (LMM stats, main effect of sleep stage: χ^2^(1)=11.75, p=6.09E-4). This effect was driven entirely by fast spindles (LMM stats, main effect of spindle type: χ^2^(1)=16.91, p=3.92E-5). Importantly, we found a significant stage x spindle type interaction (LMM stats: χ^2^(1)=12.97, p= 3.16E-4, followed by a one sample t-test for fast spindles during NREM3 vs zero: t(47.1)=5.94, p= p=0.0001), suggesting that only the density of fast spindles occurring during NREM3 was modulated by motor learning.

**Figure 2.**
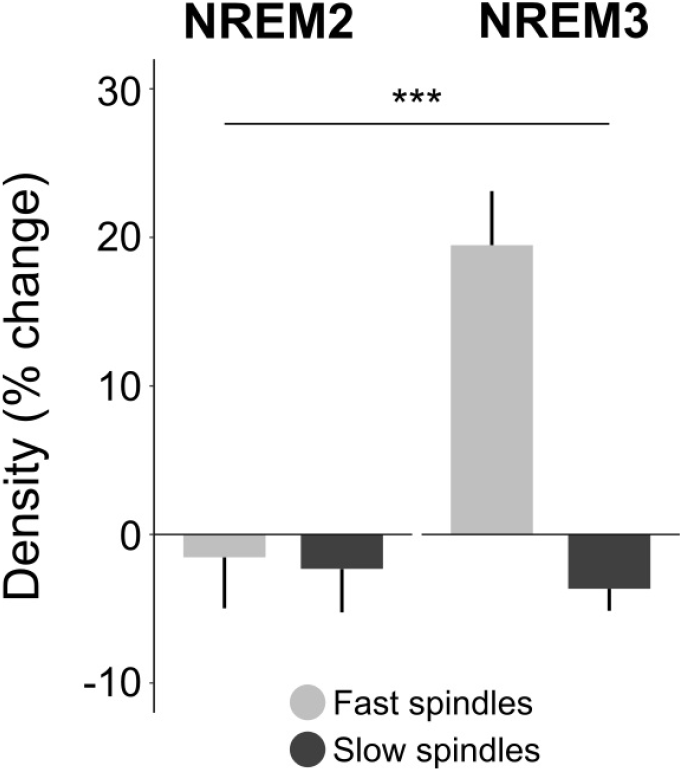
Visuomotor adaptation modulates the density of fast spindles during NREM3. The barplot depicts the relative difference in the density of fast and slow spindles (mean ± SE) between VMA and CTL sessions (VMA-CTL)/ CTL*100), during NREM2 and NREM3. Visuomotor adaptation increased the density of fast spindles during NREM3. *** p<0.001 indicates the sleep stage by spindle type interaction.

Figure 3 illustrates the topographic distribution of this metric for the VMA and CTL sessions (first two columns), as well as for the relative change between them ((VMA-CTL)/CTL*100) described in Figure 2. Note that, for both VMA and CTL sessions, fast spindles were observed over parietal and central brain regions, whereas slow spindles were rather concentrated more frontally. Specially salient was the strong modulation of VMA on the density of fast spindles (up to 36%) during NREM3 (third column), which exhibited a somewhat asymmetric pattern. Quantification of this effect between Left and Right hemispheres yielded a significant difference, with the LH showing higher density of fast spindles than the RH (relative change in density, mean ± SE: *LH=30.21±6.35 %*; *RH=14.97±5.33 %*; LMM stats: main effect of hemisphere: χ^2^(1)=5.51, p=0.02). In sum, our findings show that visuomotor adaptation increases the density of fast spindles during NREM3, in a somewhat inter-hemispheric manner.

**Figure 3.**
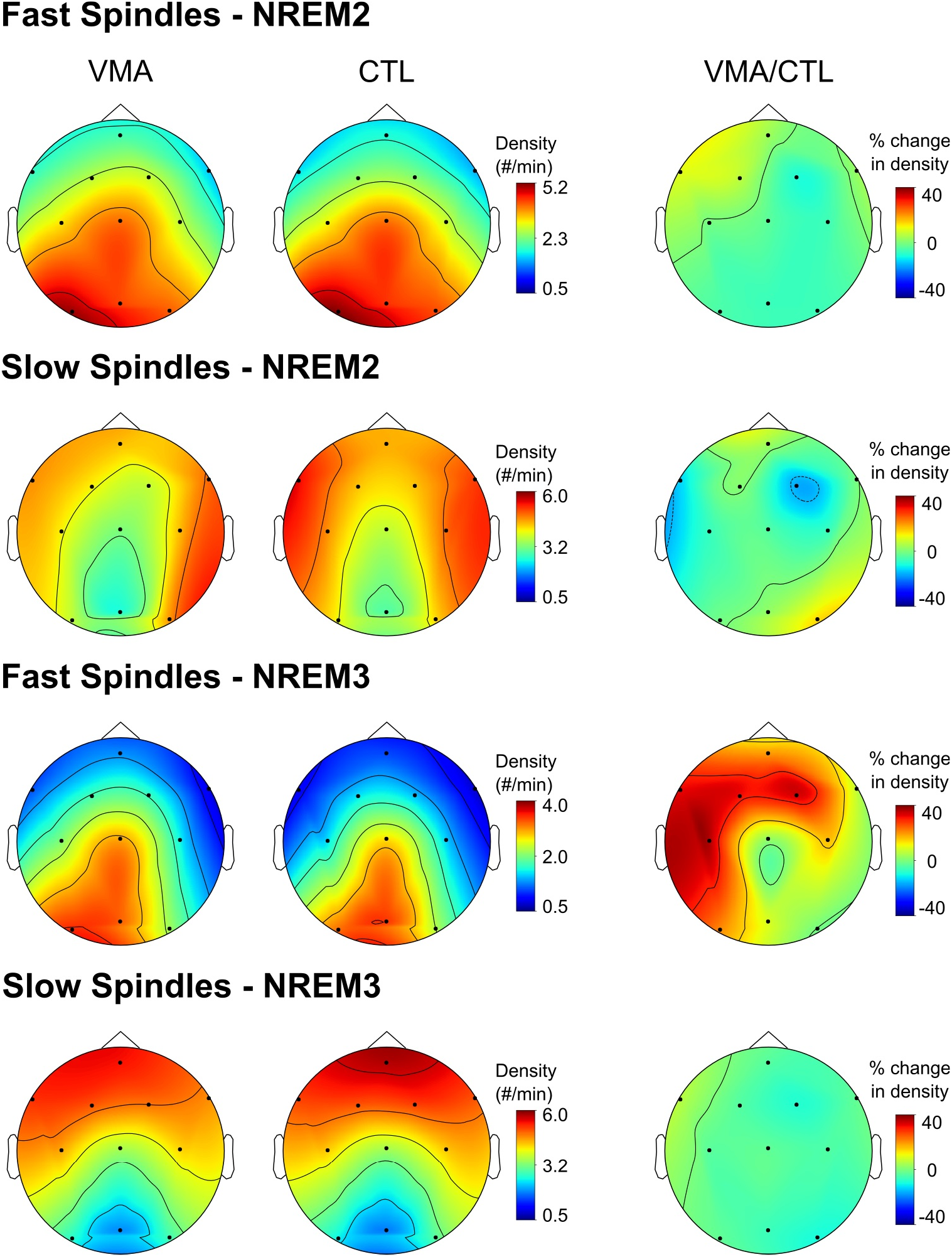
Topographic distribution of the density of fast and slow spindles during NREM2 and NREM3. Shown are the topographic plots for the density of fast and slow sleep spindles (number per minute) corresponding to the VMA session, the CTL session and their relative difference (VMA/CTL), during NREM2 (upper two rows) and NREM3 (lower two rows). Relative differences between VMA and CTL were computed according to the function (VMA-CTL)/CTL*100). Colorbars represent the density of sleep spindles for the VMA and CTL conditions, and for their percent change, respectively. Only the density of fast spindles occurring during NREM3 was modulated by motor learning (LMM stats: χ^2^(1)=12.97, p= 3.16E-4, followed by a one sample t-test for fast spindles during NREM3 vs zero: t(47.1)=5.94, p= p=0.0001)

### Effect of visuomotor adaptation on slow-wave sleep

After studying the activity of sleep spindles, we explored the impact of motor learning on slow wave oscillations. To address the main aim of our study regarding the synchrony between slow oscillations and spindles, we examined whether visuomotor adaptation modulated the density of slow oscillations (~1Hz). Given that SOs are the hallmark of NREM3 and that, as observed in the previous section, sleep spindles were mostly modulated during NREM3, we quantified SOs during this stage (Riedner et al., 2007; Mölle et al., 2011). We found that visuomotor adaptation did not increase the density of slow oscillations above that observed in the control condition (Figure 4.A; relative change in density, mean ± SE: *NREM3=−1.9 ± 0.77 %*; one sample t-test vs zero: t(9.02)=−1.07, p=0.31).

**Figure 4.**
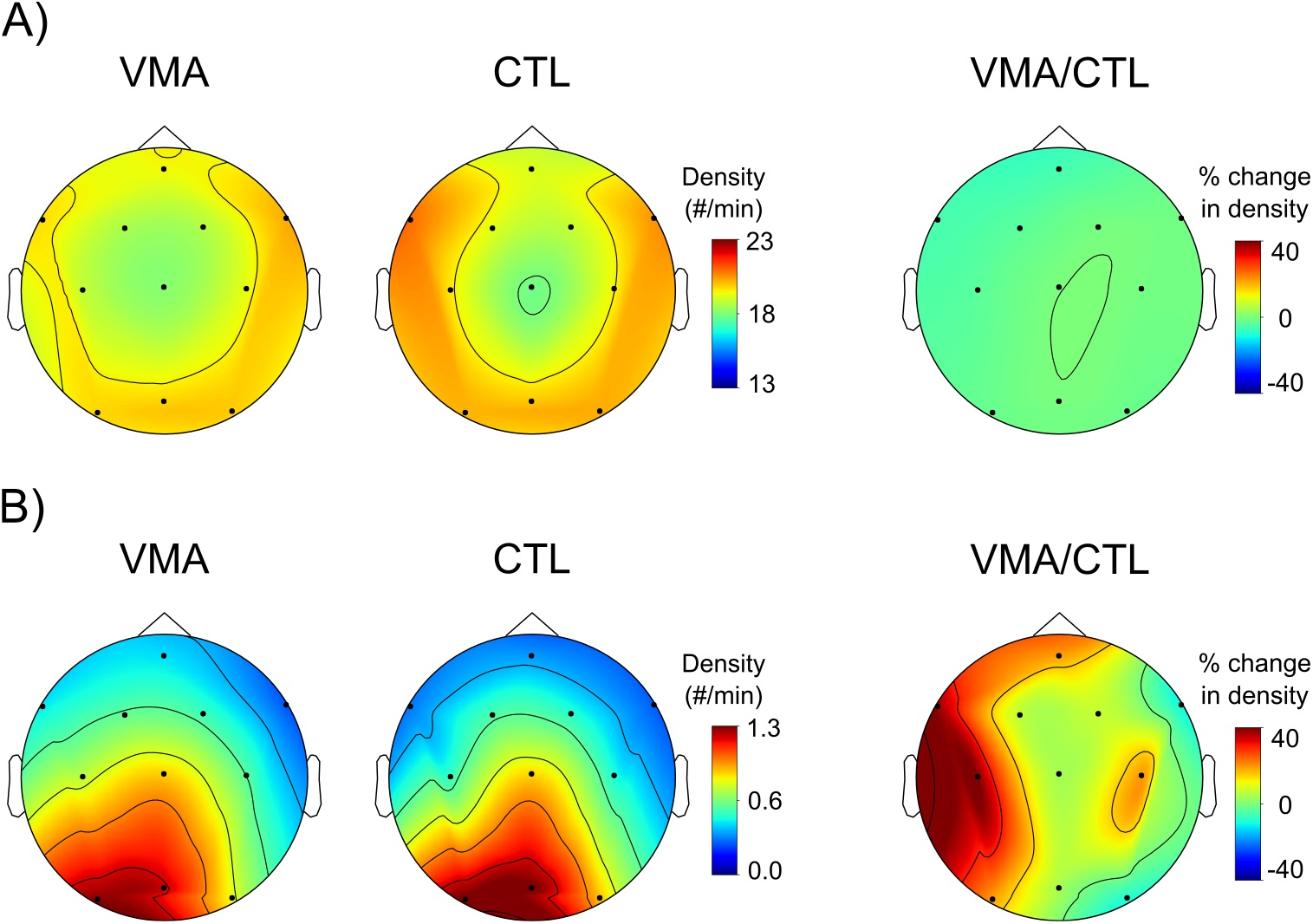
Visuomotor adaptation modulates the spindle-SO coupling during NREM3. *A) VMA did not impact on the density of slow oscillations.* Shown are the topographic plots for the density of slow oscillations for the VMA and CTL sessions and their relative difference (VMA/CTL). We found that VMA did not modulate the density of SOs during NREM3 (one sample t-test vs zero: t(9.02)=−1.07, p=0.31). *B) Local modulation of spindle-SO coupling.* Shown are the topographic plots for the density of fast spindles coupled with a slow oscillation for the VMA and CTL sessions and their relative difference (VMA/CTL). Visuomotor adaptation significantly increased the global density of spindle-SO couplings during NREM3 (one sample t-test vs zero: t(8.23)=2.62, p=0.03). Relative differences were computed according to the function (VMA-CTL)/CTL*100). Colorbars represent the density of these metrics for the VMA and CTL conditions, and their percent change, respectively.

Previous work aimed at testing the Synaptic Homeostasis Hypothesis (SHY) (Huber et al., 2004), has shown an increment in the power of delta band (1-4Hz) during the first 30 minutes of NREM sleep that decreases thereafter. Although our study was not aimed at examining changes in delta oscillations but on slow oscillations, we were interested to see if we could reproduce this result with our data. The replicability of Huber’s finding detailed in the Extended Data (Figure 4-1) suggests that SHY may be at play here, and strengthens it as a potential ubiquitous mechanism involved in the stabilization of visuomotor adaptation memories.

### Visuomotor adaptation modulates the coupling between fast spindles and slow oscillations

So far, we have shown a local modulation of motor learning on the density of fast spindles during NREM3, but not on the density of SOs. To explore the relevance of the spindle-SO coupling in the consolidation of motor memories, we quantified the amount of spindles that occurred locked to a SO during NREM3. Given that only fast spindles were modulated by sleep we examined the effect of motor learning on fast spindle-SO couplings. Even though the proportion of fast spindles coupled with a SO was similar across VMA and CTL sessions (proportion of fast spindles coupled to a SO, mean ± SE: *VMA=31.1±1.1 %, CTL=33.7±1.4 %*; LMM stats, main effect of session: χ^2^(1)=2.42, p=0.12), visuomotor adaptation significantly increased the global density of spindle-SO couplings during NREM3 relative to the control (relative change in density, mean ± SE = 17.48±5.21 %; one sample t-test vs zero: t(8.23)=2.62, p=0.03). The topographic distribution corresponding to this effect is illustrated in Figure 4.B.

Further statistical analysis on the relative difference between VMA and CTL yielded a strong inter-hemispheric modulation, suggesting that this effect was driven by the left hemisphere, i.e., the hemisphere contralateral to the trained hand (relative change in density, mean ± SE: *LH: 26.42±9.09 %*, *RH: 1.81±5.88 %*; LMM stats, main effect of hemisphere: χ^2^(1)=5.82, p=0.02).

To examine whether the increment in the fast spindle-SO coupling in fact promoted memory stabilization, we next correlated the relative change in the density of fast spindles associated with a SO during NREM3 with overnight memory retention. To establish further the specificity of this phenomenon, we contrasted this result with that obtained from correlating the relative change in the density of *uncoupled* fast spindles with memory retention. As observed in Figure 5, only fast spindles associated with a SO predicted long-term memory overnight (Pearson correlation: *Coupled fast spindles*: r1= 0.73, p=0.018 (p = 0.036 after adjusting by Bonferroni); *Uncoupled fast spindles*: r2= −0.23, p=0.53 (p = 0.99 after adjusting by Bonferroni).

**Figure 5.**
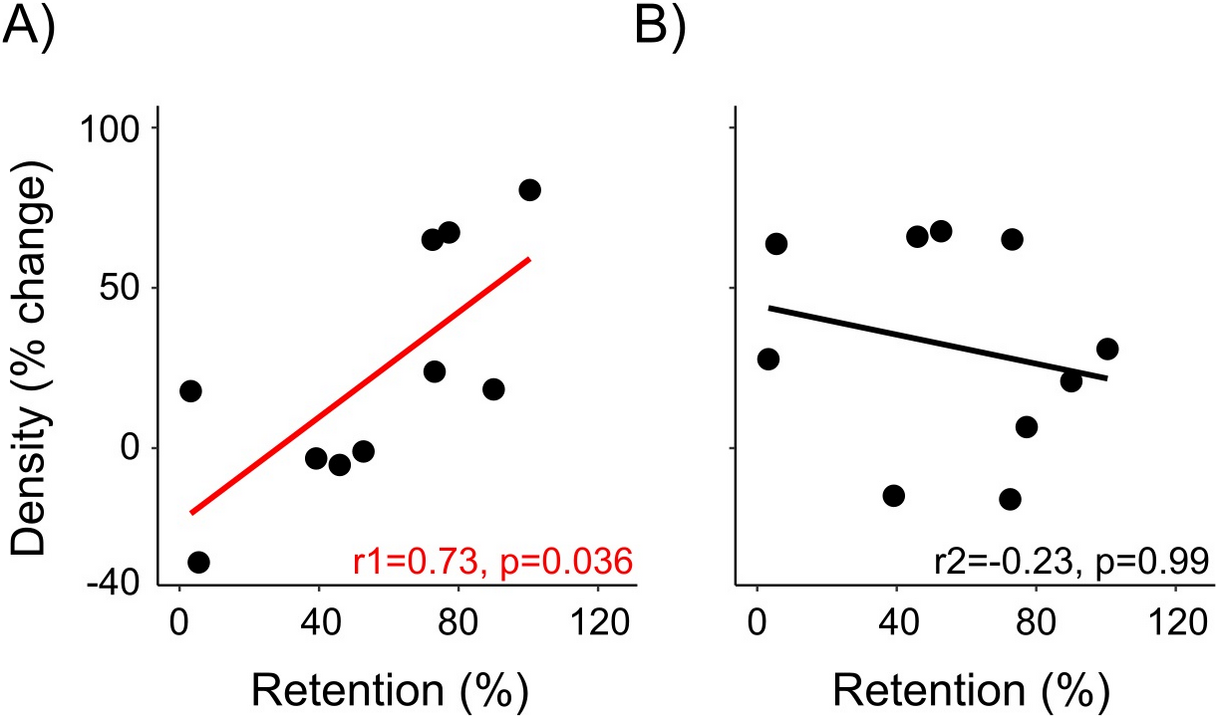
Fast spindles coupled with SOs predict motor memory stabilization. Correlation between the % change ((VMA-CTL)/CTL*100) in the density of fast spindles coupled (*A*) and uncoupled (*B*) with SOs, and overnight memory retention. Only fast spindles coupled with SOs predicted long-term memory. Shown are the dots representing the mean across all 4 electrodes of the left hemisphere for each subject, and the corresponding regression lines. Reported p-values are corrected by Bonferroni.

In sum, we found that visuomotor adaptation modulates the density of spindles but not their frequency, duration or amplitude during NREM sleep. This effect was specific to fast spindles during NREM3. Although VMA did not affect the density of SOs, it substantially increased the fast spindle-SO coupling, suggesting that motor learning may rather boost the ability of slow oscillations to promote thalamic spindles. Remarkably, adaptation modulated the degree of this coupling locally in an inter-hermispheric manner, i.e., contralaterally to the trained hand. The fact that only fast spindles coupled with a SO predicted long-term memory points to this association as a fundamental signature of motor memory consolidation.

### Phase relationship between sleep spindles and slow oscillations

Previous studies have shown that learning may influence the degree of synchrony between SOs and spindles, which is reflected in the level of grouping (more grouping is associated with less variance) of spindles around the active phase of slow oscillations (Mölle et al., 2009, 2011). To investigate whether the modulation of the fast spindle-SO coupling observed in Figure 4.B was associated with a change in the degree of synchrony, we determined the phase of the SO at which the spindle yielded its maximum peak-to-peak amplitude, and contrasted the level of spindle grouping around the mean phase for the VMA and CTL sessions.

Figure 6 illustrates a consistent phase relationship between SOs and fast spindles for both the CTL and VMA sessions (Rayleigh non-uniformity test. CTL: r=0.79, p=5e-04; VMA: r=0.96 p=1e-04). In accordance with the declarative literature, fast spindles occurred locked to the active phase of the slow oscillation, close to its positive peak (Steriade & Amzica, 1998; Amzica & Steriade, 1998; Rasch & Born, 2013; Ladenbauer et al., 2017; Helfrich et al., 2018; Muehlroth et al., 2019). We found that the level of grouping around the mean phase was similar for VMA and CTL (Wallraff’s Nonparametric Test, VMA vs. CTL: V=17, p=0.32).

**Figure 6.**
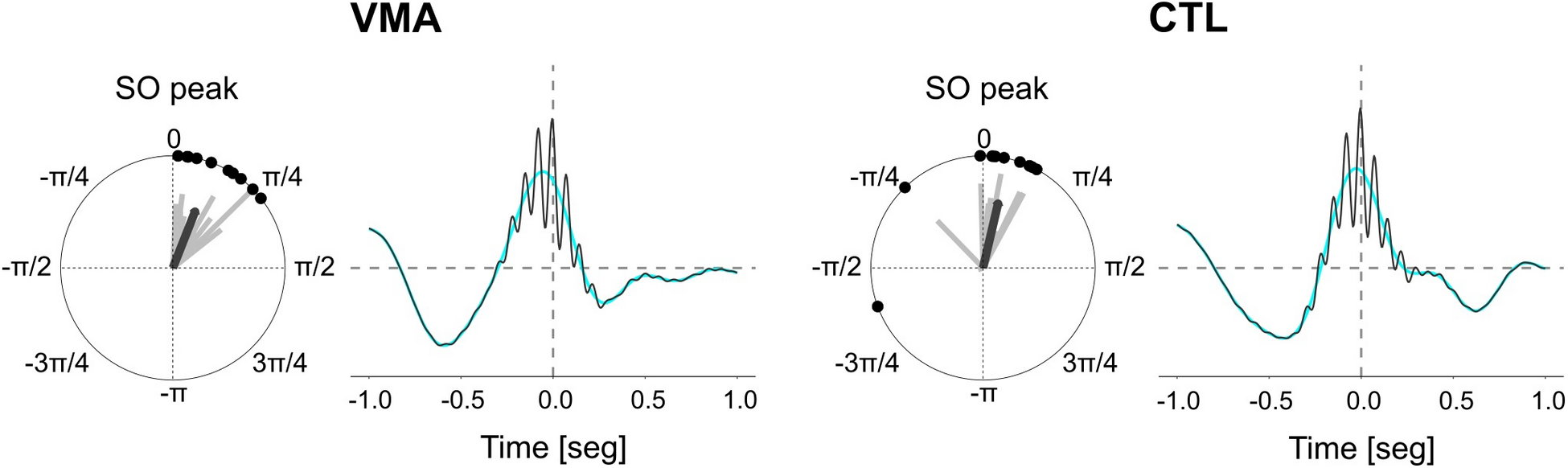
Fast Spindle-SO synchrony for VMA and CTL. Shown are the polar plots (left) and the overlaid of fast spindles onto a slow oscillation (right) for VMA and CTL. Polar plots illustrate the mean coupling phase between fast spindles and slow oscillations for each subject (gray lines/black dots) and across all subjects (black arrow), corresponding to the left hemisphere. Signal overlays depict the grand average of EEG signal for fast spindle-SO coupling events time locked to the spindle’s maximum amplitude (x =0.0), where the cyan line is the signal filtered in the slow oscillations frequency band (0.5-1.25 Hz) and the black line is the signal filtered in the fast spindles band (12-16 Hz).

In sum, we showed that fast spindles occurred mainly during the active phase of SOs independently of the condition (CTL or VMA). Even though VMA increased the coupling between slow oscillations and spindles it did not influence their level of synchrony, suggesting that this parameter may not provide relevant information regarding motor memory stabilization.

## Discussion

The degree of coupling between slow oscillations (SO) and spindles appears to be critical for sleep-dependent consolidation of declarative memories. Here, we examined whether this mechanism operates in the stabilization of motor memories. To this end, we measured the impact of learning a visuomotor adaptation task on the density of slow oscillations, spindles and their coupling during NREM sleep. Adaptation modulated both the fast spindle density and the degree of fast spindle-SO coupling locally, in an inter-hemispheric manner, during NREM3. Remarkably, the density of fast spindles associated with slow oscillations, but not uncoupled spindles, predicted memory retention overnight, underscoring the importance of this association in the stabilization of procedural memories.

Unlike motor sequence learning, the mechanisms involved in the consolidation of motor adaptation have remained somewhat elusive for over a decade. This may obey to the failure of retrograde interference protocols at unveiling a gradual recovery of the memory trace (for a comprehensive literature review refer to Krakauer et al., 2019). Using an alternative anterograde interference approach, we have recently shown evidence supporting the stabilization of visuomotor adaptation memories within a 6-hour window post training (Albert et al., 2019; Lerner et al., 2020). This time window is in line with that reported for motor sequence learning (Walker et al. 2003; Korman et al. 2007; Cantarero et al. 2013). Tracking functional connectivity during this period has allowed us to identify a network including motor, premotor and posterior parietal areas, mostly contralateral to the trained hand, that peaks about 6 hours post training and predicts long-term memory overnight (Della-Maggiore et al., 2017). Using the same experimental paradigm, here we identified a sleep related modulation of brain oscillations over these cortical areas that also predicts long-term memory overnight. This anatomical congruency is consistent with the hypothesis that sleep promotes the consolidation of memory representations in local networks that may be active during learning (e.g., Klinzing et al., 2019). The inter-hemispheric modulation of brain oscillations observed in our study is consistent across the different sleep metrics examined and, thus, stresses the specificity of our findings.

There are some major differences worth noticing regarding the neural signatures modulated by learning in our experimental paradigm compared to motor sequence learning. So far sleep consolidation of motor sequence learning has been linked to changes occurring during NREM2. For example, offline gains in performance correlate positively with the density of sleep spindles (Nishida & Walker, 2007; Boutin et al., 2018), as well as with the amount of time spent in NREM2 (Walker et al., 2002; Nishida & Walker, 2007). This is in sharp contrast with our study, in which visuomotor adaptation modulated the number of spindles exclusively during NREM3. Another important discrepancy between the two types of motor learning is the actual signal modulated by training. Overnight improvements in performance associated with motor sequence learning have been linked to an increment in the density of isolated sleep spindles whereas in our work only spindles coupled to a slow oscillation predicted long-term memory. Further work will be needed to systematically compare both motor tasks and establish whether these discrepancies reflect mechanistic differences inherent of the two types of learning, the experimental design (such as the time elapsed between training and sleep) and/or the sleep stage analyzed.

Our results show a strong association between the increase in the number of fast spindles coupled with SOs and the ability to retain information overnight. The fact that uncoupled spindles were not related to long-term memory suggests that only coupled spindles may have promoted synaptic plasticity. How may the enhanced coupling between slow oscillations and fast spindles drive motor memory stabilization? Using two-photon imaging, Niethard and collaborators (2018) found that cortical pyramidal neurons are more active, their dendrites disinhibited and, thus, more sensitive to excitatory inputs during the coupling of these oscillations than during their occurrence alone, which may facilitate dendritic plasticity. It has been proposed that it is during these events that newly encoded representations are reactivated (Steriade et al., 1998). This hypothesis finds support in a motor skill learning study carried out in rodents (Ramanathan et. al., 2015), showing task-related neural replay in close temporal association with fast spindles and slow oscillations. We speculate that a similar mechanism based on neural reactivation (Robertson & Genzel, 2020) may explain why coupled but not uncoupled spindles predicted long-term memory overnight.

The spindle-SO coupling has been proposed as a key association enabling systems consolidation of declarative memories. According to this hypothesis newly encoded memories initially stored in hippocampal and cortical networks, are reactivated during slow wave sleep and gradually integrated with existing memories at the systems level. This process is thought to depend on the close synchrony between slow oscillations, sleep spindles and hippocampal ripples (Rasch & Born, 2007; Diekelmann & Born, 2010). Another -non-exclusive- account of memory consolidation is the Synaptic Homeostasis Hypothesis (SHY), according to which synaptic weights potentiated by learning during wake are downscaled by sleep. Empirically, SHY would manifest as an increment in the power of delta oscillations during the beginning of sleep that decreases as the night progresses (Tononi & Cirelli, 2006). This process is thought to improve the signal-to-noise ratio for strongly potentiated synapses. Our results are in line with both accounts: we found that learning modulated the coupling between fast spindles and slow oscillations, as well as the power of delta early during NREM sleep. This opens the possibility that visuomotor adaptation memories may go through both these processes for their stabilization.

Finally, it is important to acknowledge that, unlike the work by Huber and collaborators (Huber et al., 2004; Landsness et al., 2009), a couple of labs have failed to find a beneficial effect of sleep on visuomotor adaptation (Donchin et el., 2002; Doyon et al., 2009). Yet, we want to emphasize that these studies showed substantial methodological differences from ours and Huber’s. For example, Donchin and collaborators used force-field adaptation, an experimental paradigm that differs from visuomotor adaptation in many parameters due to the distinct – proprioceptive-nature of the perturbation (Krakauer et al., 1999). On the other hand, Doyon and collaborators implemented large −mirror- rotations (180 degrees) that unlike the small rotation used in our study, are learned and consolidated through different mechanisms (Bedard & Sanes, 2011; Gutierrez-Garralda et. al., 2013; Telgen et. al., 2014). In fact, mirror reversal learning is associated with shifts in time-accuracy tradeoff and offline gains, features that are more common to skill learning tasks than to visuomotor adaptation tasks. Beyond the differences in the experimental paradigms, none of these studies controlled the timing between training and sleep in the way that we and Huber did (practically 10 minutes took place between the end of training and bedtime). The close proximity between training and sleep has been shown to be key in promoting a benefit of sleep for different type of material (e.g., Schönauer et al., 2015; Sawangjit et al., 2018). Another potential limitation of our study is the sample size. Although we chose a repeated measures design to improve power by reducing inter-subject variability, the sample size is not large. One of the problems of within-subjects design like ours, is the level of attrition, which increases with the number of visits to the lab. Here, subjects came to the lab four times in a month: the first time for an interview, and then three additional times for the familiarization, VMA and CTL sessions. Yet, despite the limited sample size, several factors such as the hemispheric asymmetry observed consistently across several EEG metrics, the reproducibility of Huber’s findings and the differential correlation between coupled and uncoupled spindles with long-term memory support the robustness of our results.

In conclusion, we have shown that visuomotor adaptation modulates the density and the level of coupling between fast spindles and slow oscillations in an inter-hemispheric manner during NREM3. Interestingly, only fast spindles associated with slow oscillations predicted memory retention overnight, pointing to a role of this coupling in motor memory consolidation. Our findings provide compelling evidence for a role of the spindle-SO coupling in the stabilization of motor memories, and open the possibility of a common mechanism operating at the basis of procedural and declarative memory.

## Acknowledgments

This work was supported by the Argentinian Ministry of Defence (PIDDEF), and the Argentinian Agency for the promotion of Science and Technology (FONCyT, ANPCyT).

